# Crystal structure of a mammalian CMTM6 and its interaction model with PD-L1

**DOI:** 10.1101/2023.01.03.522599

**Authors:** Senfeng Zhang, Qingrong Xie, Chunting Fu, Yongbo Luo, Ziyi Sun, Xiaoming Zhou

## Abstract

CKLF-like MARVEL transmembrane domain-containing protein 6 (CMTM6) is a master regulator of PD-L1. By binding PD-L1 at the plasma membrane and recycling endosomes, CMTM6 prevents the lysosomal degradation of PD-L1 and maintains its cell surface expression, thus stabilizing the inhibitory PD-1/PD-L1 axis. However, the mechanism of CMTM6/PD-L1 interaction is unknown. Here we report the first experimentally determined structure of CMTM6 from bovine. Combined with a low-resolution cryo-EM map, computational docking analysis and a protein binding assay an interaction model between CMTM6 and PD-L1 was proposed, providing a structural framework for the CMTM6 regulation on PD-L1.

## Main text

Immune checkpoint blockade therapy targeting PD-1/PD-L1 has been approved for treating various cancers, which highlights the important role that the inhibitory axis of PD-1/PD-L1 plays in immune tolerance(1, 2). Recent studies identify CKLF-like MARVEL transmembrane domain-containing protein 6 (CMTM6) as a master regulator of PD-L1(3, 4). By binding PD-L1 at the plasma membrane and recycling endosomes, CMTM6 maintains the cell surface expression of PD-L1 by preventing it from lysosomal targeting and degradation(3, 4). However, the mechanism of CMTM6/PD-L1 interaction is unknown, which hinders further understanding of the CMTM6 regulation on PD-L1.

To elucidate the structural basis of CMTM6/PD-L1 interaction, we first sought to determine a structure of CMTM6. To do so, 12 mammalian *CMTM6* genes, including from human and mouse, were screened for expression. Among them, the bovine CMTM6 protein (bCMTM6) displayed the best biochemical properties (Fig. 1A) and was crystallized. In the end, the bCMTM6 structure (bCMTM6_xtal_) was solved by molecular replacement using an AlphaFold-predicted model of bCMTM6 (bCMTM6_AF_, https://alphafold.com/entry/Q3ZBE8)(5). The final bCMTM6_xtal_ structure was refined to 3.50 Å with reasonable map quality (Fig. 1B and Table 1) and contains residues 33-165 of bCMTM6, and the amino-terminus (32 residues) and the carboxy-terminus (17 residues) were not modeled due to weak electron densities.

**Figure 1.**
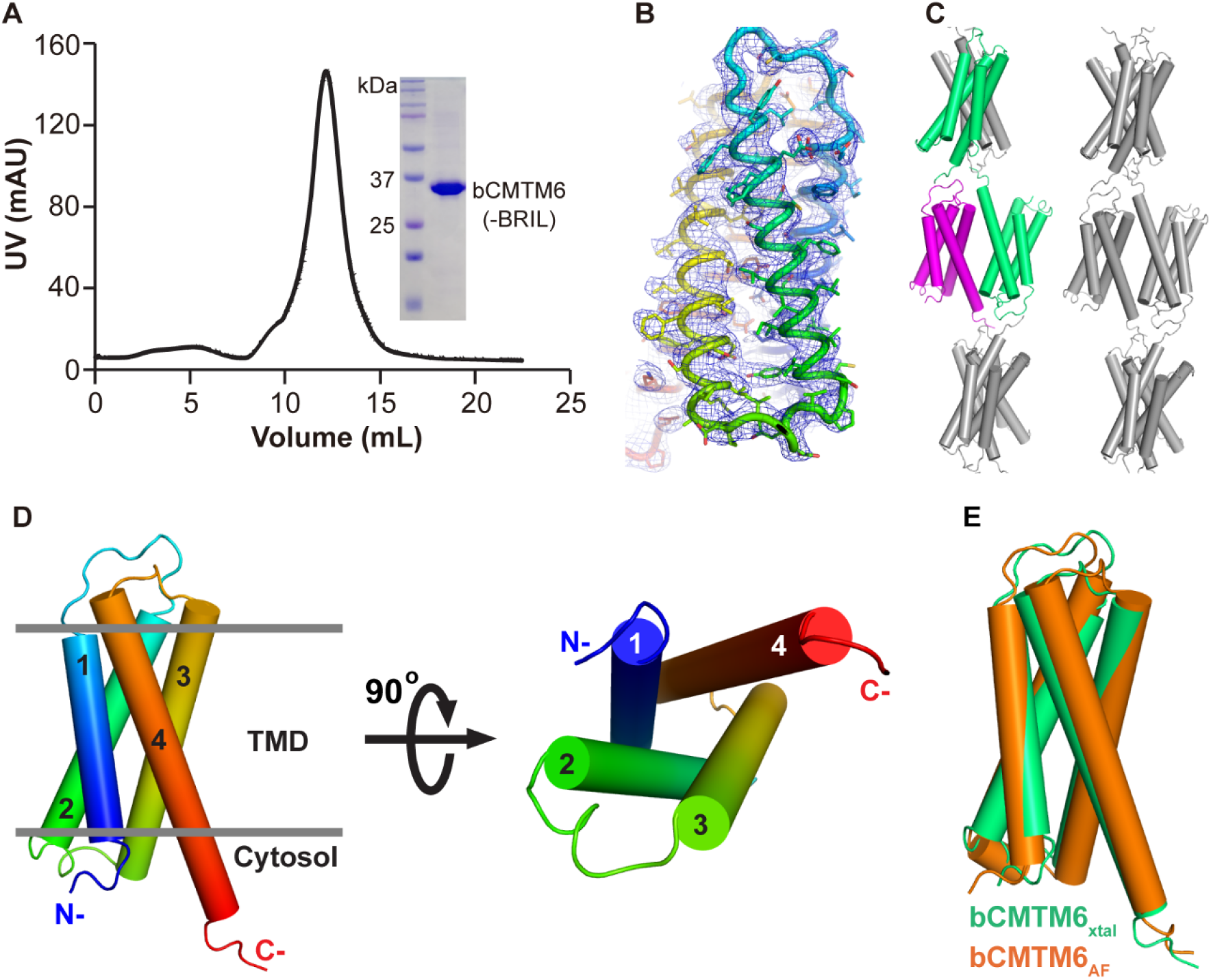
Crystal structure of bCMTM6. (A) The elution profile of bCMTM6(-BRIL) (see Extended methods for details) on a size-exclusion column. The inset shows the SDS-PAGE analysis of the purified sample. (B) The 2F_o_-F_c_ electron density map contoured at 1.2 σ level is shown as blue mesh for one protomer of the bCMTM6_xtal_ structure, which is shown as rainbow-colored tubes. (C) Crystal packing of the bCMTM6_xtal_ structure. The two green protomers are within one asymmetric unit, whereas the magenta protomer forms a crystallographic dimer with the green protomer in the center of the panel. (D) The crystal structure of one bCMTM6_xtal_ protomer is rainbow-colored from the N-to the C-terminus. Four transmembrane helices are displayed as cylinders numbered from 1 to 4. Left, viewed parallel to the membrane; Right, viewed from the cytosol. The membrane is indicated by two grey lines throughout this manuscript. (E) Superposition of the AlphaFold-predicted bCMTM6_AF_ model (in orange) onto one bCMTM6_xtal_ protomer (in green).

**Table 1.**
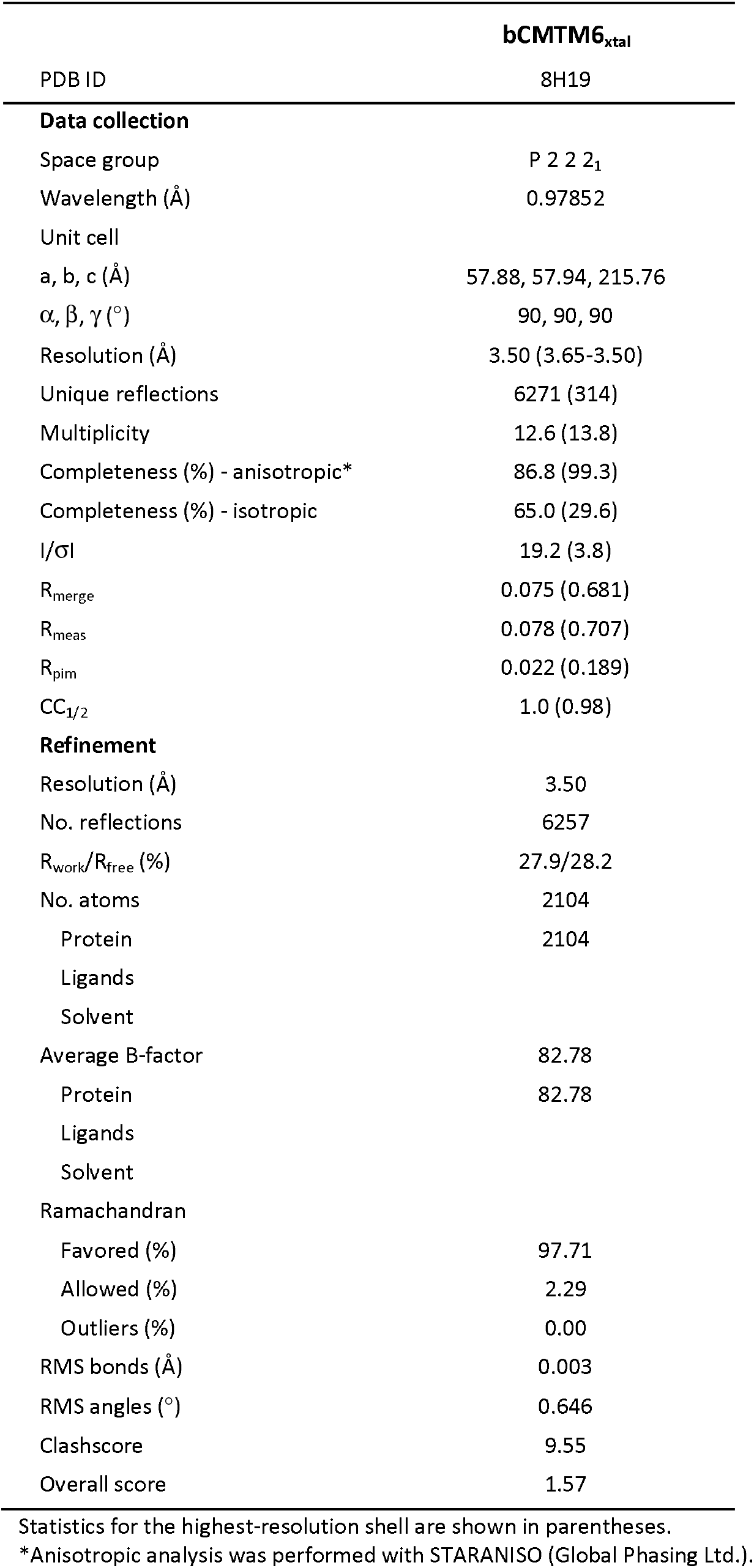
Data collection and refinement statistics.

The bCMTM6_xtal_ structure contains two protomers in its asymmetric unit (Fig. 1C). Although crystal packing analysis revealed a dimeric organization of two bCMTM6 molecules, the dimer is packed in opposite directions via the transmembrane domain (TMD) (Fig. 1C), suggesting that the crystallographic dimer of bCMTM6_xtal_ is less likely to occur in biological membranes. Therefore, structural analysis of bCMTM6_xtal_ was focused on one single protomer. Each protomer contains four transmembrane helices, with amino- and carboxy-termini both in cytosol (Fig. 1D, left panel). Viewed from the cytosolic side, the four helices of TMD are arranged counter-clockwise from TM1 to TM4 (Fig. 1D, right panel). Not surprisingly, structural alignment between bCMTM6_AF_ and a bCMTM6_xtal_ protomer yielded an all-Cα root mean square deviation (RMSD) of 1.6 Å, indicating that the two models are alike (Fig. 1E). Nonetheless, it is also noteworthy that structural details including helix positioning and loop conformations are relatively different between bCMTM6_xtal_ and bCMTM6_AF_ (Fig. 1E).

Then, the next step is to determine a bCMTM6/PD-L1 complex structure. Both CMTM6 and PD-L1 are highly conserved proteins among mammalian species(6). For example, bCMTM6 and human CMTM6 (hCMTM6) sequences are 72% identical and 87% homologous, whereas bovine PD-L1 (bPD-L1) and human PD-L1 (hPD-L1) share 73% sequence identity and 92% similarity when aligned by CLUSTALW(7). Therefore, we tested if an interaction is also preserved between bCMTM6 and hPD-L1 so that we could utilize bCMTM6 with superior biochemical properties to probe the CMTM6/PD-L1 binding mechanism with hPD-L1, which is of more biomedical significance than bPD-L1.

Interestingly, bCMTM6 and hPD-L1 displayed robust interaction using a pull-down assay (Fig. 2A and 2B, lane 4). Therefore, we attempted to solve the bCMTM6/hPD-L1 complex structure using both X-ray crystallography and cryo-EM single-particle reconstruction. Unfortunately, crystallization of the bCMTM6/hPD-L1 complex was not successful. Meanwhile, cryo-EM single-particle reconstruction yielded a low-resolution (8-9 Å) volume showing the shape of detergent micelles containing the bCMTM6/hPD-L1 complex (Fig. 2C). However, the low-resolution volume failed to reveal details about the transmembrane region of the complex, probably due to difficulties in 3D-reconstruction as the bCMTM6/hPD-L1 complex is small (∼50 kDa) and also lacks any soluble domains that are fixed to TMD (Fig. 2A)(8).

**Figure 2.**
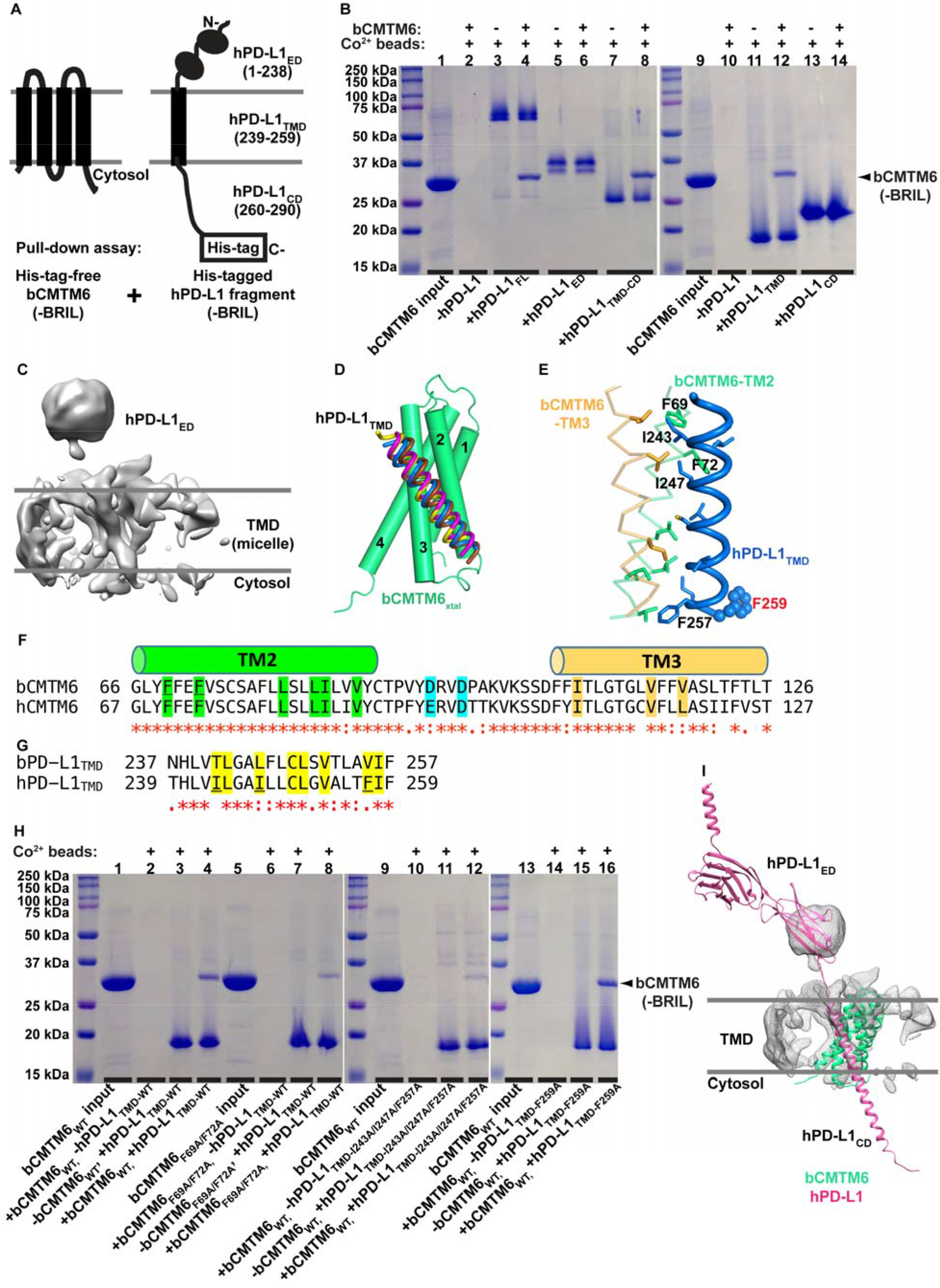
The interaction model of bCMTM6/hPD-L1. (A) A scheme showing the pull-down assay setup. For all bCMTM6 and hPD-L1 variants, a BRIL module is fused to the C-terminus of the protein of interest to improve expression (see Extended methods for details). All purified bCMTM6 variants are His-tag free, whereas all purified hPD-L1 variants are C-terminally His-tagged. (B) A pull-down experiment using purified hPD-L1 fragments to pull down purified bCMTM6 protein is examined by SDS-PAGE analysis. Signs and labels above and below the gel indicate the components present in the pull-down experiment. The position of the bCMTM6 band is indicated by an arrowhead on the right. Molecular weight of protein standards is labeled on the left. (C) A low-resolution (8.4 Å) cryo-EM volume of the bCMTM6/hPD-L1 complex in detergent micelles. (D) Four top-scored docking models of hPD-L1_TMD_ (tubes in four colors) docked into the bCMTM6_xtal_ structure (in green). (E) Hydrophobic interactions between hPD-L1_TMD_ (blue tube) and bCMTM6’s TM2 (green ribbon) and TM3 (light orange ribbon) in the top-scored docking model. Residues expected to participate in the hydrophobic interactions are rendered as sticks. A non-participating residue (F259 of hPD-L1_TMD_) is displayed as spheres. (F)-(G) Sequence alignment between part of bCMTM6 and hCMTM6 (F), and between bPD-L1_TMD_ and hPD-L1_TMD_ (G) by CLUSTALW(7). Residues of hydrophobic interactions are highlighted in green (CMTM6-TM2), light orange (CMTM6-TM3) and yellow (PD-L1_TMD_). Two acidic residues in the CMTM6-TM2/TM3 loop are highlighted in cyan. F69/F72 of bCMTM6-TM2, and I243/I247/F257 of hPD-L1_TMD_ are underscored. (H) A pull-down experiment using purified hPD-L1_TMD_ variants to pull down purified bCMTM6 variants as indicated, similar to panel (B). (I) A bCMTM6_xtal_/hPD-L1_full-length_ model fitted in the low-resolution cryo-EM density of panel (C). Of note, only the nearest domain of hPD-L1_ED_ to the membrane is visible in the cryo-EM volume.

To further explore the binding mechanism between bCMTM6 and hPD-L1, we utilized computational docking analysis and a pull-down binding assay (Fig. 2A). Since hPD-L1 is a single-pass transmembrane protein containing an extracellular domain (ED, residues 1-238), a TMD (residues 239-259) and a cytosolic domain (CD, residues 260-290) (Fig. 2A)(4), we first tested which domain(s) of hPD-L1 contributes to its binding to bCMTM6. Intriguingly, hPD-L1_TMD-CD_ and hPD-L1_TMD_ bound robustly to bCMTM6 (Fig. 2B, lanes 8 and 12), while hPD-L1_ED_ or hPD-L1_CD_ showed no detectable binding (Fig. 2B, lanes 6 and 14), indicating that hPD-L1_TMD_ is mainly responsible for the bCMTM6 interaction. This result is largely consistent with a previous study showing that the PD-L1 TMD is required for efficient interaction with CMTM6(4).

Since hPD-L1_TMD_ is a well-folded transmembrane helix and bCMTM6_xtal_ is an experimentally determined structure, computational docking with both reliable structures is expected to yield more confident results. We then created docking models through the HDOCK web server(9) using a bCMTM6_xtal_ protomer as the receptor structure and an AlphaFold-predicted hPD-L1_TMD_ model (https://alphafold.com/entry/Q9NZQ7, residues 239-259) as the ligand structure (see Extended methods for details). Interestingly, the top-scored four models all have the hPD-L1_TMD_ helix clustered in the TM2/TM3 face of the bCMTM6_xtal_ protomer (Fig. 2D), suggesting that the interaction between bCMTM6-TM2/TM3 and hPD-L1_TMD_ may possibly represent a real binding pattern between bCMTM6 and hPD-L1. Using the top-scored bCMTM6_xtal_/hPD-L1_TMD_ model for analysis, the interaction between bCMTM6-TM2/TM3 and hPD-L1_TMD_ is primarily mediated by hydrophobic residues (Fig. 2E). In bCMTM6, the participating residues include Phe69, Phe72, Leu80, Leu83, Ile84 and Val87 from bCMTM6-TM2, and Val101, Ile108, Val115 and Val118 from bCMTM6-TM3, which are extremely conserved in hCMTM6 except that Val118 of bCMTM6 has a corresponding Leu119 in hCMTM6 (Fig. 2F). In hPD-L1_TMD_, the participating residues are Ile243, Leu244, Ile247, Cys250, Leu251, Val253, Phe257 and Ile258, which are also very conserved in bPD-L1 (Fig. 2G). To validate the binding pattern between bCMTM6-TM2/TM3 and hPD-L1_TMD_, we generated a bCMTM6-TM2/TM3 double mutant (bCMTM6_F69A/F72A_) and an hPD-L1_TMD_ triple mutant (hPD-L1_TMD-I243A/I247A/F257A_), and tested their ability to bind their wild-type partners. Consistently, both bCMTM6_F69A/F72A_ and hPD-L1_TMD-I243A/I247A/F257A_ caused apparently reduced binding compared to the wild-type pair (Fig. 2H, lanes 8 and 12 compared to lane 4), supporting the binding pattern proposed above. Meanwhile, binding to wild-type bCMTM6 remained unaffected in an hPD-L1_TMD_ mutant at residue 259 (hPD-L1_TMD-F259A_, Fig. 2H, lane 16), which is not in contact with bCMTM6 according to the bCMTM6_xtal_/hPD-L1_TMD_ model (Fig. 2E) and served as a negative control.

For discussion purposes, we also created a bCMTM6_xtal_/hPD-L1_full-length_ model by aligning the AlphaFold-predicted model of full-length hPD-L1 (https://alphafold.com/entry/Q9NZQ7) to the bCMTM6_xtal_/hPD-L1_TMD_ model through matching the hPD-L1_TMD_ portion (Fig. 2I). Interestingly, the bCMTM6_xtal_/hPD-L1_full-length_ model could be fitted readily into the low-resolution cryo-EM volume of the bCMTM6/hPD-L1 complex (Fig. 2I), suggesting that the bCMTM6_xtal_/hPD-L1_full-length_ model may be somewhat plausible. Of note, CD of the AlphaFold-predicted hPD-L1_full-length_ model is mainly α-helical (Fig. 2I), whereas an NMR study suggests that hPD-L1_CD_ is disordered and its basic residues interact with acidic membrane lipids to regulate its ubiquitination(10). Further study is required to investigate if PD-L1_CD_ assumes different conformational states to fulfill its function. Meanwhile, it has been suggested that CMTM6 may prevent PD-L1 from ubiquitination, likely through interactions with PD-L1_CD_(4). According to the bCMTM6_xtal_/hPD-L1_full-length_ model (Fig. 2I), it is tempting to postulate that the basic residues of hPD-L1_CD_ interact with the acidic residues of the bCMTM6-TM2/TM3 loop (Asp94 and Asp97), which are conserved in hCMTM6 (Glu95 and Asp98) (Fig. 2F), thus protecting hPD-L1_CD_ from being ubiquitinated. Therefore, it will be desirable to experimentally determine a CMTM6/PD-L1 complex structure in future studies to further elucidate the CMTM6 regulation on PD-L1.

## Acknowledgements and funding sources

Diffraction data used in this study were collected on the beamline BL19U1 of National Facility for Protein Science in Shanghai (NFPS) at Shanghai Synchrotron Radiation Facility (SSRF). The authors thank the beamline staff for assistance during data collection. Cryo-EM data were collected at SKLB West China Cryo-EM Center and were processed at the Duyu High Performance Computing Center of Sichuan University. This work was supported in part by Oversea Scholar Science and Technology Project by Sichuan Human Resources and Social Security Department to ZS, and the National Natural Science Foundation of China (NSFC) grant 31770783, and the 1.3.5 Project for Disciplines of Excellence grant ZYYC20014 by West China Hospital of Sichuan University to XZ.

## Data availability

The atomic coordinates and structure factors of the bCMTM6_xtal_ structure have been deposited in the Protein Data Bank under the accession code 8H19.

## Author contributions

XZ and ZS conceived and supervised the project, and wrote the manuscript with input from all authors. SZ performed structural and functional studies; QX performed functional studies; CF performed structural studies; YL collected cryo-EM data; and all authors analyzed the data.

## References

1. He X & Xu C (2020) Immune checkpoint signaling and cancer immunotherapy. Cell Res 30(8):660–669.

2. Han Y, Liu D, & Li L (2020) PD-1/PD-L1 pathway: current researches in cancer. Am J Cancer Res 10(3):727–742.

3. Burr ML, et al. (2017) CMTM6 maintains the expression of PD-L1 and regulates antitumour immunity. Nature 549(7670):101–105.

4. Mezzadra R, et al. (2017) Identification of CMTM6 and CMTM4 as PD-L1 protein regulators. Nature 549(7670):106–110.

5. Jumper J, et al. (2021) Highly accurate protein structure prediction with AlphaFold. Nature 596(7873):583–589.

6. Takeuchi H, et al. (2020) Expression Analysis of Canine CMTM6 and CMTM4 as Potential Regulators of the PD-L1 Protein in Canine Cancers. Front Vet Sci 7:330.

7. Thompson JD, Higgins DG, & Gibson TJ (1994) CLUSTAL W: improving the sensitivity of progressive multiple sequence alignment through sequence weighting, position-specific gap penalties and weight matrix choice. Nucleic Acids Res 22(22):4673–4680.

8. Nygaard R, Kim J, & Mancia F (2020) Cryo-electron microscopy analysis of small membrane proteins. Curr Opin Struct Biol 64:26–33.

9. Yan Y, Tao H, He J, & Huang SY (2020) The HDOCK server for integrated protein-protein docking. Nat Protoc 15(5):1829–1852.

10. Wen M, et al. (2021) PD-L1 degradation is regulated by electrostatic membrane association of its cytoplasmic domain. Nat Commun 12(1):5106.

