## Extended methods for "Crystal structure of a mammalian CMTM6 and its interaction model with PD-L1"

**Supplementary information**

**Extended methods**

**Protein expression and purification.** The gene encoding the full-length wild-type bovine CMTM6 (bCMTM6, NCBI accession code: NP_001030238.1) and human PD-L1 (hPD-L1, NCBI accession code: NP_054862.1) were synthesized by Genewiz (Suzhou, China) in a modified pPICZ plasmid (Thermo Fisher Scientific) containing a carboxy-terminal tag of Tobacco etch virus (TEV) protease recognition site and decahistidine. The bacterial cytochrome b562RIL (BRIL)(1) was fused to the carboxy-end of the bCMTM6 and hPD-L1 proteins before the TEV site to improve their expression and stability. All site-directed mutants and hPD-L1 fragments were generated by polymerase chain reaction (PCR) and were verified by DNA sequencing, and were subsequently cloned into the same vector as bCMTM6/hPD-L1. Proteins of interest were expressed and purified according to a previously published procedure(2) with slight modifications. Briefly, the plasmid containing the sequence of bCMTM6(-BRIL) fusion protein was linearized and transformed into yeast (*Pichia pastoris* strain GS115) by lithium chloride/single-stranded carrier DNA/polyethylene glycol method according to manufacturer's manual (Thermo Fisher Scientific). The bCMTM6(-BRIL) protein was overexpressed by adding 1% (v/v) methanol and 2.5% (v/v) dimethyl sulfoxide (DMSO) at $OD600_{nm}$ of 3~5 and shaking at 30 °C for 24 h. Cell pellets were resuspended in lysis solution (LS) containing 20 mM Tris-HCl pH 7.5, 150 mM NaCl, 10% (v/v) glycerol, 1 mM phenylmethanesulfonyl fluoride (PMSF) and 2 mM β-mercaptoethanol, and were lysed by an ATS AH-1500 high-pressure homogenizer (Shanghai, China) at 1,300 MPa. Undisrupted cells and cell debris were separated by centrifugation at 3,900 x g for 20 min at 4°C, and membranes were collected by ultracentrifugation at 130,000 x g for 1 h at 4°C. Protein was extracted by addition of 1% (w/v) n-dodecyl-β-D-maltopyranoside (DDM, Anatrace) and 0.1% (w/v) cholesteryl hemisuccinate (CHS, Anatrace) at 4 °C for 2 h and the extraction mixture was centrifuged at 200,000 x g for 25 min at 4 °C. The supernatant was incubated with $Co^{2+}$ resin in the presence of 20 mM imidazole pH 8.0 at 4 °C for 1 h, then the mixture was loaded onto a cobalt metal affinity column, washed with 20 bed-volume of LS containing 1 mM DDM, 0.01% (w/v) CHS and 40 mM imidazole pH 8.0, and eluted with LS supplemented with 1 mM DDM, 0.01% (w/v) CHS and 250 mM imidazole pH 8.0. Expression and purification of hPD-L1(-BRIL) and fragments followed the same protocol as bCMTM6(-BRIL), except for hPD-$L1_{ED}$(-BRIL) (residues 1-238) and hPD-$L1_{CD}$(-BRIL) (residues 260-290), which are soluble proteins and their purification omits the membrane harvest step and does not use detergents.

**Crystallization.** Affinity-purified bCMTM6(-BRIL) protein was concentrated to ~5 mg/ml and was treated with $\alpha$-chymotrypsin (TLCK-treated, Sigma-Aldrich) at a 1:50 ratio (chymotrypsin: bCMTM6, w/w) for 9 min at 18 °C to generate a stable core. The digestion was stopped by 10 mM PMSF. The mixture was loaded onto a Superdex 200 Increase 10/300 GL column (Cytiva Life Sciences) equilibrated in 20 mM Tris-HCl pH7.5, 150 mM NaCl, 5 mM β-mercaptoethanol, 13 mM n-nonyl-beta-D-glucoside (NG) and 0.04% (w/v) CHS and was further purified by size-exclusion chromatography (SEC). SEC-purified bCMTM6 was then concentrated to 8-10 mg/ml as approximated by ultraviolet absorbance, and 500 nl of protein solution was mixed with an equal volume of crystallization solution manually in a vapor diffusion sitting-drop setup and was incubated at 18 °C. The bCMTM6 crystals grew in 0.5 M KCl, 0.05 M sodium citrate pH 6.0, 16% (v/v) PEG 400, 0.1 M L-proline and 10 mM $K_2Pt(CN)_4$, which usually appear within four weeks, and reach full-size in six weeks. The bCMTM6 crystals were cryo-protected by raising the precipitant (PEG 400) concentration to final 36% (v/v) with a 2% incremental step, and were flash-frozen in liquid nitrogen.

**X-ray data collection and structure solution.** X-ray diffraction data were collected on the beamline BL19U1(3) of National Facility for Protein Science in Shanghai (NFPS) at Shanghai Synchrotron Radiation Facility (SSRF). The data were indexed, integrated, and scaled using the autoPROC pipeline package (Global Phasing Ltd.)(4), which includes XDS(5), AIMLESS (CCP4 package)(6) and anisotropy analysis by STARANISO (Global Phasing Ltd.). The bCMTM6 structure (bCMTM6$_{xtal}$) was solved by molecular replacement using an AlphaFold-predicted model of bCMTM6 (bCMTM6$_{AF}$, https://alphafold.com/entry/Q3ZBE8)(7). Manual model building and refinement was carried out using Coot(8) and phenix.refine(9), and Molprobity(10) was used to monitor and improve protein geometry. Non-crystallographic symmetry (NCS) restraints were applied throughout the refinement to improve maps. The data collection and refinement statistics were generated using phenix.table_one(9) combined with the output from STARANISO analysis (Global Phasing Ltd.), and were summarized in Table 1. All structural figures and RMSD calculations were performed in PyMOL (Schrödinger, LLC) and UCSF Chimera(11).

**Pull-down assay.** For pull-down analysis, affinity-purified bCMTM6(-BRIL) variants were treated with TEV protease to remove the His-tag, while affinity-purified hPD-L1(-BRIL) variants (and fragments) kept their His-tag, and all proteins were further purified by SEC. In a pull-down experiment, SEC-purified hPD-L1 variants were first incubated with $Co^{2+}$ beads for 40 min at 4 °C, followed by addition of His-tag-free bCMTM6 variants to continue incubation for 1 h at 4 °C in the presence of 30 mM imidazole pH 8.0. The $Co^{2+}$ beads were collected by centrifugation at 5,000 x g for 1 min and washed twice with the assay buffer before being analyzed by SDS-PAGE gels.

**Cryo-EM sample preparation and image acquisition.** The bCMTM6(-BRIL)/hPD-L1(-BRIL) complex was obtained by the pull-down method described above. Briefly, affinity-purified His-tagged hPD-L1(-BRIL) was used to pull down His-tag-free bCMTM6(-BRIL), and the complex was further purified by SEC in 20 mM Tris-HCl pH 7.5, 150 mM NaCl, 5 mM β-mercaptoethanol and 0.5 mM DDM. Freshly SEC-purified bCMTM6/hPD-L1 complex was then concentrated to ~10 mg/ml, and 3 µl of the protein solution was applied to glow-discharged (45 s) Quantifoil R2/1 300-mesh gold grids (Quantifoil Micro Tools GmbH, Germany). The grids were blotted with standard Vitrobot filter paper for 2.5 s at 4 °C under 100% humidity and plunged into liquid ethane using a Vitrobot Mark IV (Thermo Fisher Scientific). The frozen grids were loaded into a Titan Krios electron microscope (Thermo Fisher Scientific) operated at 300 kV with the condenser lens aperture at 50 µm. Microscope magnification was at 165,000× (0.85 Å per pixel). Movie stacks were collected automatically using the EPU software (Version 2.9.0.1519REL, Thermo Fisher Scientific) on a K2 Summit direct electron camera (Gatan) equipped with a Quantum GIF energy filter with an energy slit of 20 eV in counting mode. Data were collected at 5 raw frames per second for 6 s, yielding 30 frames per stack and a total exposure dose of 63.3 e-/$Å^2$. A total of 4,669 movie stacks were collected at a defocus range of -1.0 to -1.8 µm.

**Cryo-EM data processing and 3D reconstruction.** The movie stacks of the bCMTM6/hPD-L1 complex were motion-corrected and dose-weighted using the MotionCor2 program(12), and the rest of the data processing was carried out in cryoSPARC 4.0(13). A criterion of "CTF fit resolution < 4 Å" excluded 628 movie stacks. Blob picking was first carried out using 200 micrographs that picked 57,175 particles, which were used for an initial 2D classification. Class averages representing projections of the bCMTM6/hPD-L1 complex were selected as templates for automated particle picking from the full dataset. A total of 656,095 particles were picked from 4,041 micrographs. Particles extracted from the dataset were subjected to reference-free 2D classification, and particles in good 2D classes with a visible hPD-L1$_{ED}$ domain in addition to the main micelle body were selected (407,147 good particles in total) for an initial 3D model reconstruction (Ab-Initio, 3 classes). The Ab-Initio results were then fed into a Heterogeneous refinement with three initial volumes. The Ab-Initio reconstruction and Heterogeneous refinement steps were performed in six parallel runs, and particles of the best Heterogeneous refinement class with visible hPD-L1$_{ED}$ from each run were combined and duplicate particles were removed, resulting in 164,874 particles. These particles were then subjected to a final three-class Heterogeneous refinement, and the best 3D class was selected, containing 84,296 particles. Unfortunately, a non-uniform refinement(14) with these particles yielded a 3D volume of ~8.4 Å, which only shows the shape of detergent micelles containing the bCMTM6/hPD-L1 complex with very limited details offered (see Fig. 2C). Since the low-resolution density was not sufficient to solve the structure of the bCMTM6/hPD-L1 complex, it was not deposited in the Protein Data Bank or the Electron Microscopy Data Bank.

**Docking of hPD-L1$_{TMD}$ in bCMTM6$_{xtal}$.** Computational docking was performed using the HDOCK web server(15) (http://hdock.phys.hust.edu.cn/). For the input of receptor, the bCMTM6$_{xtal}$ structure was prepared as a .pdb file containing only one protomer. For the input of ligand, an AlphaFold-predicted hPD-L1$_{TMD}$ model (https://alphafold.com/entry/Q9NZQ7, residues 239-259) was also prepared as a .pdb file. Docking was performed in the template-free docking mode. Docked models were scored and ranked automatically by HDOCK, and were also manually examined to remove incorrectly placed hPD-L1$_{TMD}$ (e.g. being placed outside the membrane).

### Supplementary references


1. Chu R*, et al.* (2002) Redesign of a four-helix bundle protein by phage display coupled with proteolysis and structural characterization by NMR and X-ray crystallography. *J Mol Biol* 323(2):253-262.
2. Meng F, Xiao Y, Ji Y, Sun Z, & Zhou X (2022) An open-like conformation of the sigma-1 receptor reveals its ligand entry pathway. *Nat Commun* 13(1):1267.

3. Zhang WZ*, et al.* (2019) The protein complex crystallography beamline (BL19U1) at the Shanghai Synchrotron Radiation Facility. *NUCL SCI TECH* 30:170-181.
4. Vonrhein C*, et al.* (2011) Data processing and analysis with the autoPROC toolbox. *Acta Crystallogr D Biol Crystallogr* 67(Pt 4):293-302.
5. Kabsch W (2010) Xds. *Acta Crystallogr D Biol Crystallogr* 66(Pt 2):125-132.
6. Winn MD*, et al.* (2011) Overview of the CCP4 suite and current developments. *Acta Crystallogr D Biol Crystallogr* 67(Pt 4):235-242.
7. Jumper J*, et al.* (2021) Highly accurate protein structure prediction with AlphaFold. *Nature* 596(7873):583-589.
8. Emsley P & Cowtan K (2004) Coot: model-building tools for molecular graphics. *Acta Crystallogr D Biol Crystallogr* 60(Pt 12 Pt 1):2126-2132.
9. Afonine PV*, et al.* (2012) Towards automated crystallographic structure refinement with phenix.refine. *Acta Crystallogr D Biol Crystallogr* 68(Pt 4):352-367.
10. Davis IW*, et al.* (2007) MolProbity: all-atom contacts and structure validation for proteins and nucleic acids. *Nucleic Acids Res* 35(Web Server issue):W375-383.
11. Pettersen EF*, et al.* (2004) UCSF Chimera--a visualization system for exploratory research and analysis. *J Comput Chem* 25(13):1605-1612.
12. Zheng SQ*, et al.* (2017) MotionCor2: anisotropic correction of beam-induced motion for improved cryo-electron microscopy. *Nat Methods* 14(4):331-332.
13. Punjani A, Rubinstein JL, Fleet DJ, & Brubaker MA (2017) cryoSPARC: algorithms for rapid unsupervised cryo-EM structure determination. *Nat Methods* 14(3):290-296.
14. Punjani A, Zhang H, & Fleet DJ (2020) Non-uniform refinement: adaptive regularization improves single-particle cryo-EM reconstruction. *Nat Methods* 17(12):1214-1221.
15. Yan Y, Tao H, He J, & Huang SY (2020) The HDOCK server for integrated protein-protein docking. *Nat Protoc* 15(5):1829-1852.
